# The relationship between distortion product otoacoustic emissions and audiometric thresholds in the extended high-frequency range

**DOI:** 10.1101/2024.07.05.601801

**Authors:** Samantha N. Hauser, Alexandra R. Hustedt-Mai, Anna Wichlinski, Hari M. Bharadwaj

## Abstract

Distortion product otoacoustic emissions (DPOAEs) and behavioral audiometry are routinely used for hearing screening and assessment. These measures provide related information about hearing status as both are sensitive to cochlear pathologies. However, DPOAE testing is quicker and does not require a behavioral response. Despite these practical advantages, DPOAE testing is often limited to screening only low and mid- frequencies. Variation in ear canal acoustics across ears and probe placements has resulted in less reliable measurements of DPOAEs near 4 kHz and above where standing waves commonly occur. Stimulus calibration in forward pressure level and responses in emitted pressure level can reduce measurement variability. Using these calibrations, this study assessed the correlation between audiometry and DPOAEs in the extended high frequencies where stimulus calibrations and responses are most susceptible to the effect of standing waves. Behavioral thresholds and DPOAE amplitudes were negatively correlated, and DPOAE amplitudes in emitted pressure level accounted for twice as much variance as amplitudes in sound pressure level. Both measures were correlated with age. These data show that with appropriate calibration methods, extended high-frequency DPOAEs are sensitive to differences in audiometric thresholds and highlight the need to consider calibration techniques in clinical and research applications of DPOAEs.

## I. INTRODUCTION

The extended high-frequency region at the base of the cochlea appears highly susceptible to dysfunction. Factors commonly associated with sensorineural hearing loss, such as aging and noise exposure, often affect hearing in the extended high frequencies. For example, many previous studies have shown that extended high-frequency hearing begins to decline at around 30 years of age, gradually worsening and extending to lower frequencies with each decade of life (Carcagno and Plack, 2020; Lee et al., 2012; Wang et al., 2021). Some studies have also shown that a history of noise exposure is associated with poorer hearing sensitivity in the extended high frequencies (Liberman et al., 2016), though not all studies of young, noise-exposed adults have replicated this finding (Bharadwaj et al., 2022). In addition to age and noise exposure, ototoxic medications and some systemic illnesses can result in elevated extended high-frequency thresholds before audiometric thresholds worsen at lower frequencies (Lough and Plack, 2022). A history of ear infections has also been linked to extended high-frequency hearing loss, even after active disease has resolved (Hunter et al., 1996). In addition, extended high-frequency hearing loss is seen in pediatric populations with otherwise normal hearing sensitivity. In a study of over 500 children aged 7 to 19, approximately 7% showed extended high-frequency hearing loss (Mishra et al., 2022).

Good extended high-frequency hearing sensitivity is less critical than the mid-frequencies for understanding of speech in quiet but likely has important implications for hearing in difficult listening conditions. Speech recognition in noise has been shown to be poorer when the signal is lowpass filtered at 8 kHz (Motlagh Zadeh et al., 2019). Extended high frequencies are also salient cues for localization (Langendijk and Bronkhorst, 1999) and loss of extended high-frequency hearing is a significant predictor of self-reported hearing difficulty in noise (Hunter et al., 2020).

Despite the prevalence of extended high-frequency hearing loss and its implications for hearing in noise, hearing thresholds are not commonly tested at those frequencies in a standard audiological assessment. Widespread clinical use of extended high-frequency audiometric testing is limited in part because it requires specialized headphones with additional calibrations that not all clinics may have. Extended high-frequency testing also prolongs testing time without a direct impact on hearing aid fitting. Even if an audiologist has the clinical time and resources to measure extended high-frequency thresholds, behavioral audiometry at any frequency requires patient feedback. Some patient populations, such as infants and young children, are unable to complete a traditional hearing testing no matter the frequency range being tested. Thus, alternative test procedures are needed.

Otoacoustic emissions (OAEs) have the potential to address these drawbacks of behavioral testing. OAEs are physiological responses to sound generated in the cochlea and measurable in the ear canal. In a healthy ear, interaction between two tones on the basilar membrane results in distortion products. The cubic distortion product (2f1-f2) can be measured by a microphone in the ear canal. The active process of the outer hair cells give rise to the cochlear non-linearities that creates this response, so absent or reduced DPOAEs are typically indicative of abnormal outer hair cell function. Because both the stimulus and the response must travel through the outer and middle ear systems, these measurements can also be affected by conductive hearing loss. So, DPOAEs must be interpreted in the context of other assessments of outer and middle ear health (e.g., otoscopy, tympanometry).

DPOAEs are widely used in the audiology clinic as a quick, non-invasive, measure of cochlear health. They are commonly used for hearing screenings in newborns and in populations that are difficult to test behaviorally (Lonsbury-Martin and Martin, 1990). DPOAEs are often more affected by noise (Lapsley Miller & Marshall, 2007) and ototoxic drugs (Stavroulaki et al., 2001) than behavioral audiometry. High frequency DPOAEs are commonly part of ototoxic monitoring programs given their high sensitivity to ototoxic damage (Reavis et al., 2008). In research settings,

DPOAEs are frequently used to monitor hearing in animal models and to assess the effects of noise, ototoxic drugs, or aging (Whitehead et al., 1992).

Though DPOAEs and the audiogram are both sensitive to hearing loss and outer hair cell dysfunction, previous comparisons between DPOAE amplitudes and audiometry have not found a one-to-one correlation between the two measurements (Gorga et al., 1997). Though part of the discrepancy between measurements results from differences in the physiological processes they reflect, extraneous factors related to the acoustic properties of DPOAEs and its measurement also contribute (Heitmann et al., 1996).

DPOAEs are generated by a combination of two basic mechanisms: nonlinear distortion and linear reflection (Shera and Guinan, 1999). Interactions between the linear reflection and non-linear distortion components of the emission results in peaks and valleys of the response. This fine structure is clearly visible when using swept tone stimuli, but it is less readily apparent when using discrete tones (Shaffer et al., 2003). Because the two components of the emission have different phase properties, the fine structure can be mitigated by measuring swept DPOAEs with an appropriate analysis window to effectively smooth the response and provide more accurate estimate DPOAE amplitudes (Abdala et al., 2015). Clinical DPOAE measurements at discrete frequencies do not control for this source of error.

Not only can interactions between the reflection and distortion components of the response alter estimates of DPOAE amplitudes, but calibration-related inaccuracies of stimuli levels have made it difficult to obtain repeatable measurements of high frequency otoacoustic emissions (Heitmann et al., 1996). In clinical otoacoustic emission systems, ear canal acoustics are assessed by an in-ear calibration prior to testing. A calibration stimulus is played, and the sound levels in the ear are measured by the microphone in the probe used to record the emissions in clinical systems. The sound level at the probe is taken as a near approximation of the sound level at the tympanic membrane. While this estimation holds for low- and mid-frequencies, the difference between levels measured at the entrance of the ear canal and at the tympanic membrane can vary significantly for higher frequencies. At the probe, sound levels result from the sum of both the forward-traveling sound wave and the waves reflected by the tympanic membrane, creating standing waves at some frequencies. The resulting valleys in the frequency response underestimate the sound levels reaching the eardrum and stimulus levels would be incorrectly adjusted to compensate. The specific frequencies affected, and the degree to which this affects the sound level reaching the eardrum, vary with probe placement and individual ear anatomy (Charaziak and Shera, 2017; Maxim et al., 2019). Variation across tests—both during repeated measurements of the same ear and across ears—can lead to as much as 20 dB differences in the level of the stimulus that reaches the tympanic membrane (Maxim et al., 2019; Scheperle et al., 2008). Calibration-related artifacts have been shown to affect DPOAEs in multiple studies (e.g., Dreisbach and Siegel, 2001, Siegel and Hirohata, 1994a, Lee et al.2012), most notably with high-frequency stimuli.

Although the inaccuracies resulting from standing waves in ear-level calibrations have long been acknowledged, the implementation of a practical solution is still relatively new for otoacoustic emissions. One such technique estimates the Thevenin-equivalent impedance and sound pressure from the probe (Scheperle et al., 2011) to separate the forward-traveling and reflected waves in the ear canal and estimate their levels independently, reducing the effect of standing waves. When using this technique, stimulus levels are measured in dB Forward Pressure Level (FPL). Separating the forward- and backward-traveling waves also makes calculation of the emissions sound level leaving the ear straightforward, so the level of the emission can be converted from sound pressure level at the microphone to dB Emitted Pressure Level (EPL) which approximates the level of sound at the tympanic membrane traveling toward the microphone. Measuring stimuli levels in FPL and the responses in EPL has been shown to decrease test-retest variability of DPOAE amplitudes compared to traditional calibrations, particularly in the high frequencies (Scheperle et al., 2008).

Given the clinical value of testing in the extended high-frequencies, and the benefits of DPOAEs over traditional behavioral audiometry, we investigated whether FPL and EPL calibration methods that increase reliability of DPOAE measurements improve DPOAE estimates of behavioral thresholds in adults. We also explored the effect of age on extended high-frequency measurements as both otoacoustic emissions and behavioral thresholds in the standard hearing range are known to covary with age (Lee et al., 2012; Oeken et al., 2000).

## II. MATERIALS AND METHODS

### A. Participants

This study included 166 participants (61 male), aged 18 to 60 years (mean = 32.3 years, SD = 13.4 years) from Purdue University and the Greater Lafayette, Indiana region. They were recruited as part of a larger study of individual differences in physiological markers of cochlear synaptopathy in the normal hearing population (Bharadwaj et al., 2022). All participants had normal hearing sensitivity in at least one ear from 250-8000 Hz, defined as thresholds less than or equal to 25 dB HL. All subjects reported no history of neurological disorders or ear pathology. All procedures were approved by the Purdue IRB (#1609018209), and participants were compensated for their time.

### B. Behavioral Audiometry

All testing took place in an electrically shielded, sound-attenuating booth. Otoscopy confirmed ear canals were clear of obstructing cerumen. As normal hearing sensitivity in the standard clinical range was a requirement for the broader study, all participants completed behavioral audiometry before further testing. Audiometry was completed using the GSI Audio Star Pro (Eden Prairie, MN) with Sennheiser HDA 200 high-frequency circumaural headphones. Thresholds at 0.25, 0.5, 1, 2, 3, 4, 6, 8, 10, 12.5, 14 and 16 kHz were determined using pulsed pure tones and the modified Hughson-Westlake procedure. Participants were asked to report detection of a tone by pushing a button. Low-frequency (0.25–2 kHz), high-frequency (3–8 kHz), and extended high-frequency (9–16 kHz) pure tone averages were calculated for each subject.

### C. Compensation for Individual Ear Canal Acoustics

As in Bharadwaj et al. (2022), the Thevenin-equivalent source pressure and impedance across frequency of the probe was estimated with the built-in ER-10X calibrator (Interacoustics). The probe was coupled to loads where the impedance can be calculated (i.e., an 8 mm diameter brass tube with five length settings). A “calibration error” was derived based on the deviation between the measured and calculated load impedance of each length of tube (Scheperle et al., 2011). Typical errors for the data collected in this study were less than 0.05. Errors of less than 1 were considered good calibrations and required before moving to ear calibration

Following probe calibration, the probe was coupled to an appropriately sized ear tip and placed securely in the subject’s ear canal. Repeated clicks were presented to estimate immittance properties of the subject’s ear and derive the appropriate voltage to FPL transfer function for a given probe fit. Low-frequency absorbance was required to be less than 29% and the admittance phase was greater than 44° to ensure the probe was adequately sealed in the ear canal before DPOAE measurements were made. DPOAE levels were converted from sound pressure level (dB SPL) to emitted pressure level (dB EPL) using the individualized transfer function derived from the participants’ in-ear calibration (Scheperle et al., 2011).

### D. Distortion Product Otoacoustic Emissions

Participants watched a captioned and muted movie of their choosing while seated comfortably in the sound-treated booth. Following in-ear calibration of the probe, primary tones were swept downward logarithmically at a rate of one-third octave/second with a frequency ratio of f1=f2/1.225. Two stimulus sweeps with overlap in frequencies were presented. For the first, f2 ranged from 16 to 4 kHz; for the second, f2 ranged from 8 to 2 kHz. We found excellent agreement in DPOAE amplitudes from the two sweeps in the overlapping frequency range. The frequencies of the primary tones were held constant for 0.5 seconds at the beginning and end of each sweep. Stimuli were presented at f1=66 and f2=56 dB FPL. Twenty-five trials of each sweep were presented. The first trial and all trials with energy greater than two times the mean absolute deviation above the median were not included in the final averaged response. A least-squares approach was used to calculate the magnitude of the primary distortion product at 2f1 - f2 after averaging the response across trials with a 0.5 second windows to extract only the distortion-component of the emission (Abdala et al., 2015). As with audiometry, DPOAE levels were summarized by averaging emissions in the high (3-8 kHz) and extended high (9–16 kHz) frequencies.

## III. RESULTS

Individual differences in ear canal acoustics were greatest at frequencies near 4 kHz and above (Figure 1) with peaks in variability around 4 and 8 kHz. These regions of highest variance correspond to the quarter-wave and half-wave resonance frequencies of the typical human ear canal. As the subjects had normal hearing sensitivity in the standard clinical frequency range (0.25-8 kHz), most participants were expected to have present DPOAEs in the high-frequency range (3-8 kHz). Our results were consistent with this expectation—high-frequency DPOAEs were above the noise floor for all 166 subjects, albeit spanning a wide range of amplitudes (-18.86 to 13.62 dB EPL). Hearing thresholds were not controlled in the extended high-frequency range, hence larger range of DPOAE amplitudes was observed for these frequencies (-36.53 to 1.64 dB EPL).

**FIG. 1.**
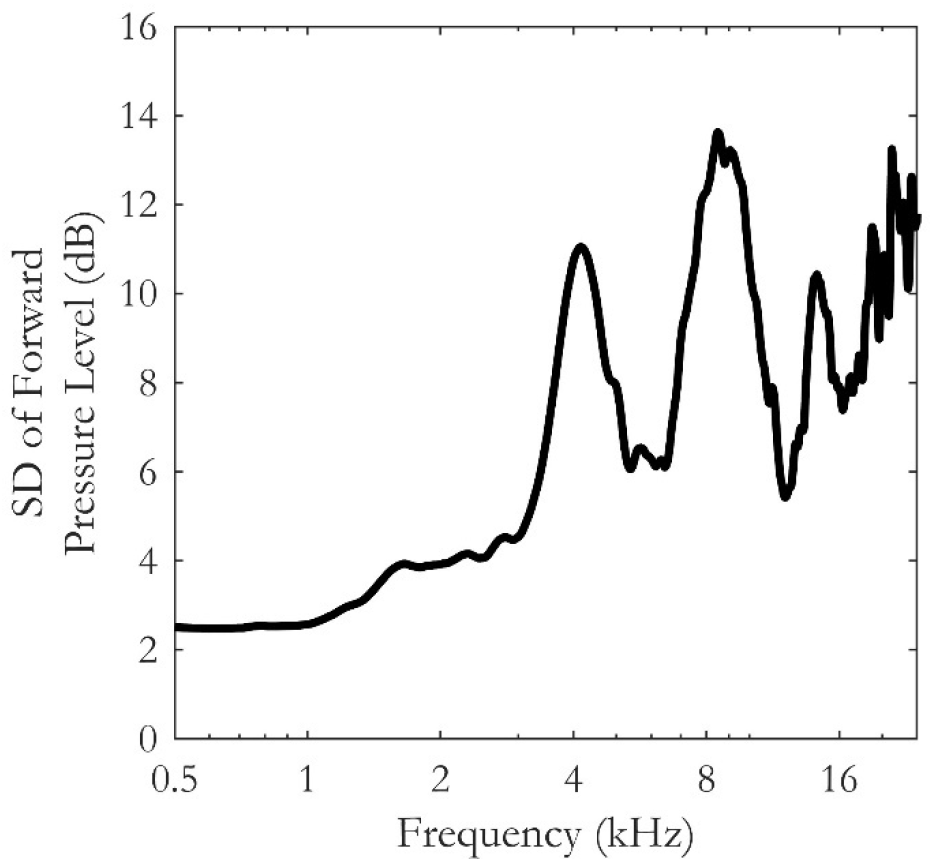
Standard deviation (SD) of in-ear calibrated forward pressure level with constant voltage stimulation (n = 166). Standard deviation increases with increasing frequency with local maxima near 4 and 8 kHz.

### A. DPOAE Relationship to Audiometry

In the extended high-frequency range, DPOAE amplitudes were negatively correlated with audiometric thresholds. Significant correlations between DPOAE amplitudes and audiometric thresholds were present whether the emission was measured in SPL (Figure 2A; r = -0.326, p<0.001) or EPL (Figure 2B; r = -0.466, p<0.001). However, the correlation when using EPL was stronger and accounted for roughly twice as much variance, though the difference was not statistically significant (Fisher’s r-to-z transform, z=-1.5, p = 0.067). Since all DPOAEs were collected only with FPL-calibrated stimuli, this dataset only allowed analysis of the effect of converting DPOAE amplitude from SPL to EPL units.

**FIG. 2.**
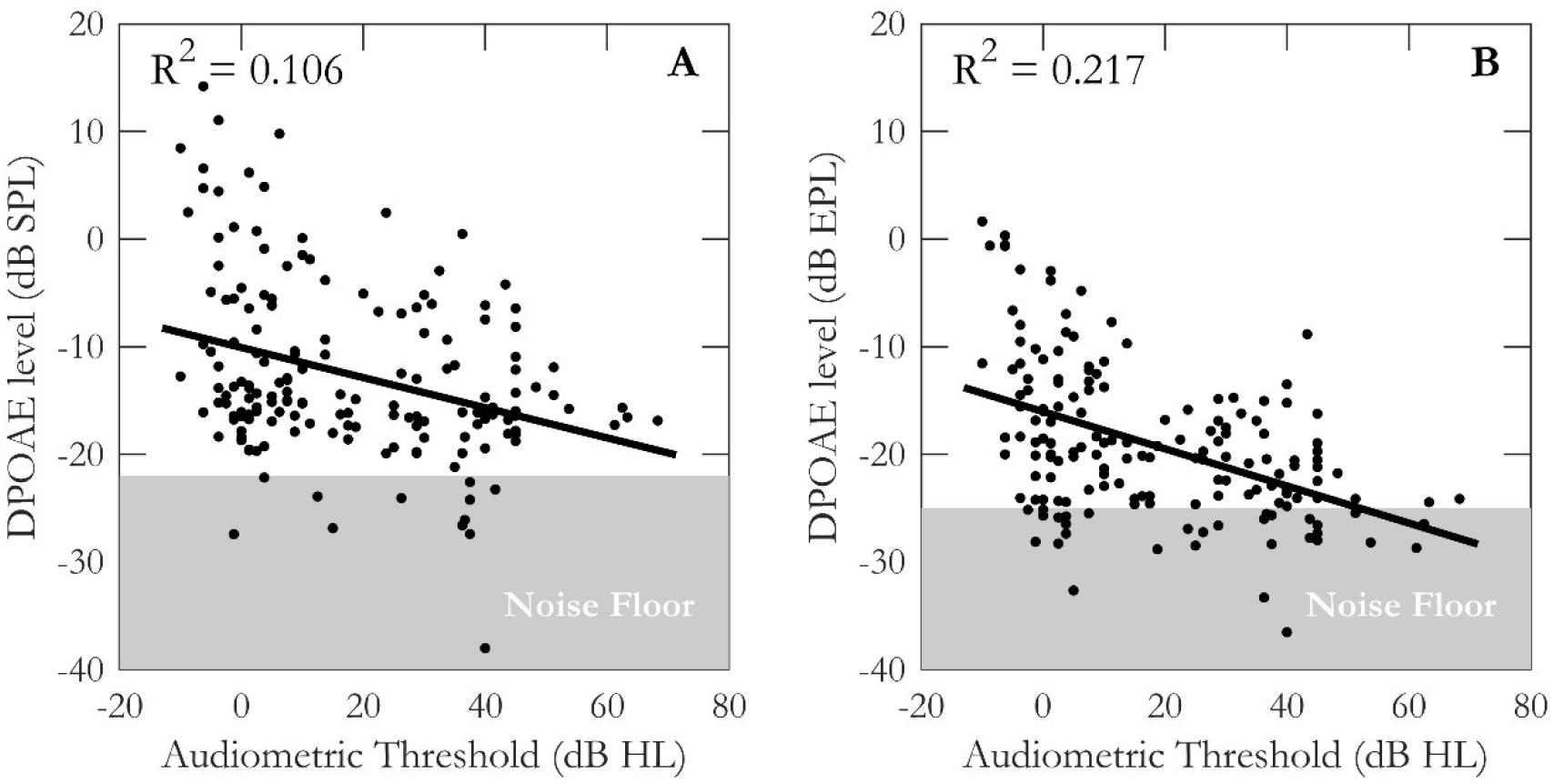
Audiometric thresholds averaged from 9-16 kHz compared to averaged DPOAE amplitudes in the same frequency range (n=166). The average noise floor for all participants’ DPOAEs is shown in grey. (A) shows the relationship when DPOAE amplitude is in Sound Pressure Level (EPL) units and (B) shows the relationship when DPOAE amplitude is in Emitted Pressure Level (SPL) units.

Although the EPL calibration method reduced variability in the correlation between DPOAE amplitudes and audiometric thresholds, DPOAE amplitudes varied among individuals, even when behavioral hearing thresholds were similar. For example, participants with extended high-frequency audiometric thresholds near zero had DPOAE amplitudes spanning a 30 dB range. Most participants with normal behavioral hearing thresholds (< 25 dB HL) in the extended high-frequency range had DPOAEs above the noise floor.

We also investigated whether high-frequency DPOAEs were predictive of extended high-frequency hearing loss, as prior studies have found (Lough and Plack, 2022). To see whether DPOAE amplitudes in one frequency range were indicative of change in higher frequency regions, we compared extended high-frequency audiometry to high-frequency DPOAE amplitudes (Figure 3). DPOAE amplitudes in the high-frequency range proved to be lower as extended high-frequency hearing thresholds worsened (r = -0.277, p<0.001). Consistent with these prior studies, our results indicate that DPOAEs from 3-8 kHz may be sensitive to extended high-frequency hearing loss and thus serve as an early indicator of hearing loss in this extended high-frequency range.

**FIG. 3.**
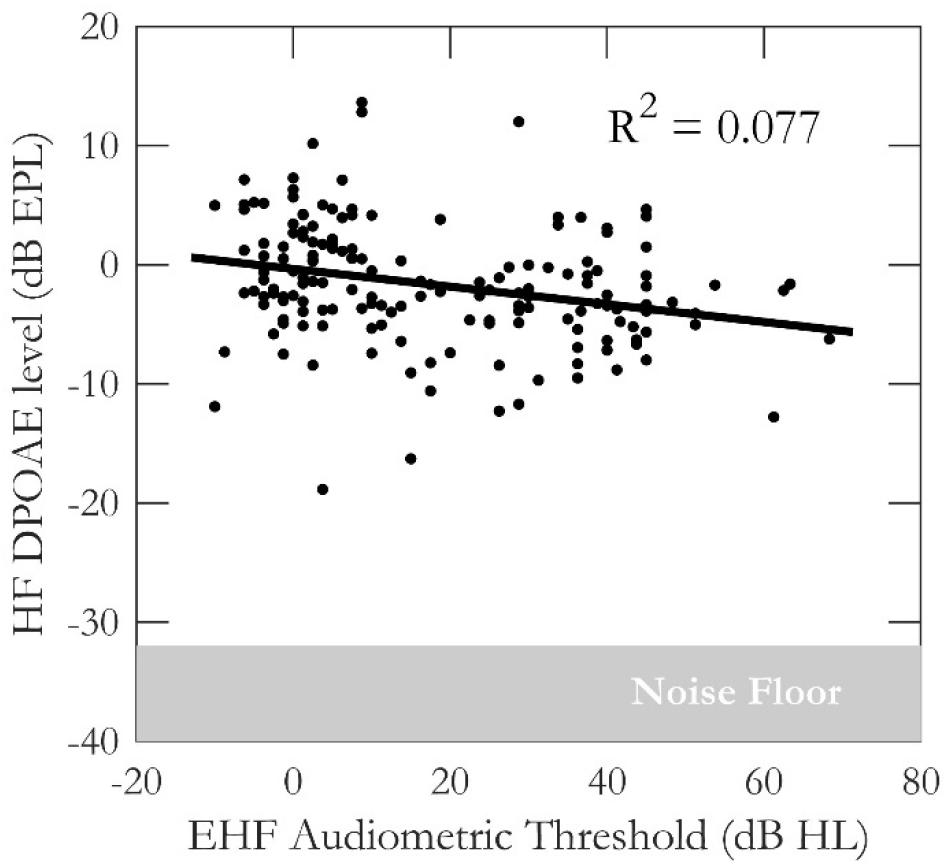
Lower high frequency DPOAE amplitudes are correlated with poorer extended high-frequency audiometric thresholds.

### B. Correlations with Age

As expected, age was significantly correlated with extended high-frequency audiometric thresholds (Figure 4A; r = 0.852, p<0.001). Audiometric thresholds in the extended high-frequency range worsened with age, despite all subjects having normal hearing through 8 kHz. In our cohort, nearly every participant over the age of 40 had elevated extended high-frequency thresholds.

**FIG. 4.**
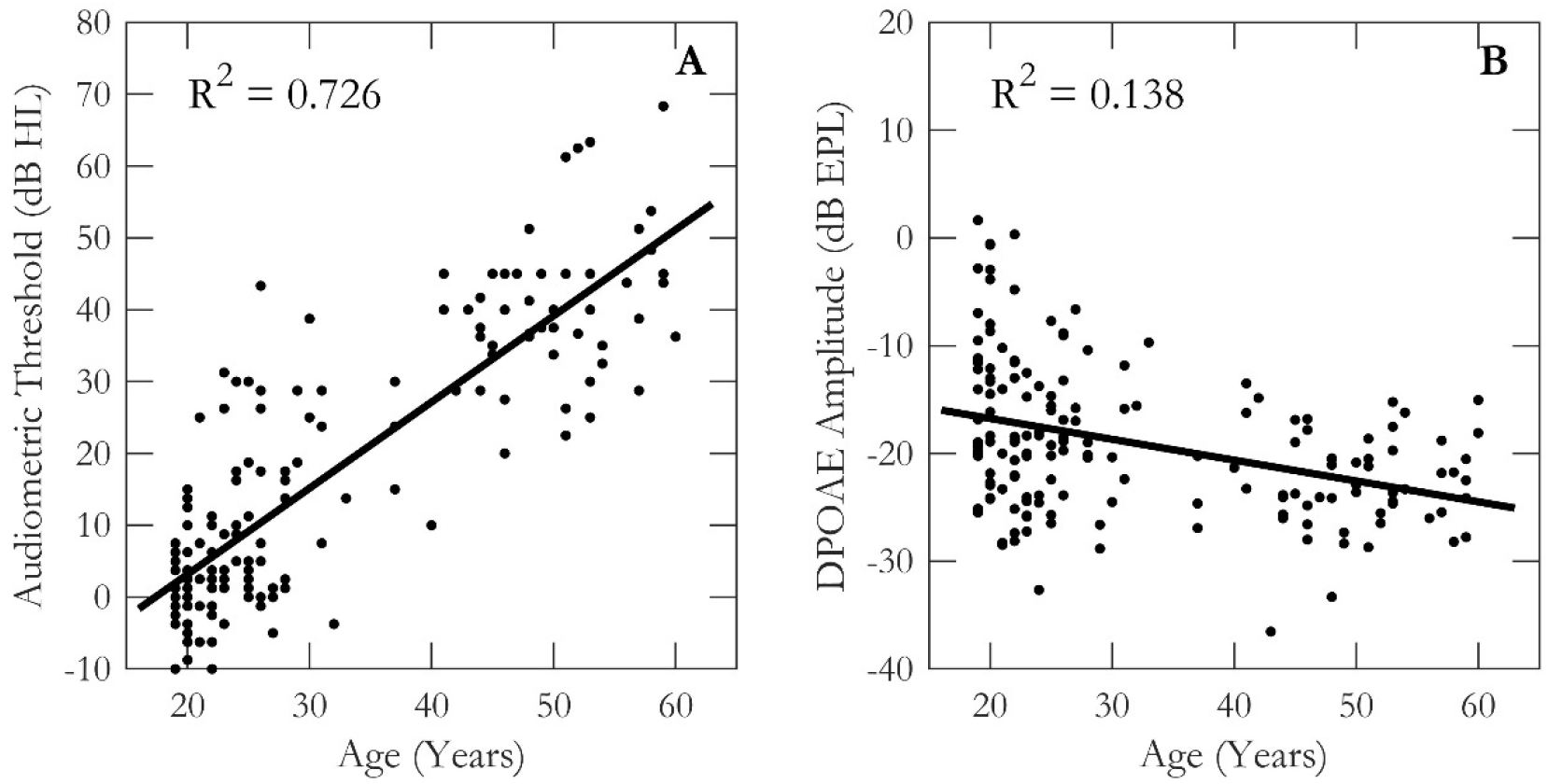
Age is correlated with both audiometric thresholds (A) and DPOAE amplitudes (B) in the extended high-frequency range.

Similarly, as age increased, extended high-frequency DPOAE amplitudes declined significantly (r = - 0.371, p<0.001).

## IV. DISCUSSION

Change in extended high-frequency thresholds is often an early indicator of cochlear pathology, but hearing in this range is not commonly assessed in the audiology clinic due to equipment and time limitations. If the methodological limitations that introduce extraneous sources of variability obscuring the relationship between DPOAE amplitudes and audiometric thresholds are overcome, DPOAEs would be an ideal tool to assess extended high-frequency hearing as they can quickly be measured without a behavioral response from the listener.

Here, we tested whether FPL and EPL calibration methods, which account for individual variation in stimulus levels at the eardrum, improve our ability to use DPOAEs to estimate behavioral thresholds in the extended high-frequency range. Our results demonstrated a strong correlation between audiometric thresholds and DPOAE amplitudes in the extended high-frequency range. This correlation was further strengthened by converting DPOAE amplitudes from SPL to

EPL units. Furthermore, we found that both behavioral thresholds and DPOAE amplitudes were highly correlated with age in the extended high-frequency range, even when hearing was normal at frequencies from 250 through 8000 Hz.

### A. Methodological Considerations for DPOAEs

Extended high-frequency testing is especially sensitive to the acoustics of the ear canal, and this has made DPOAE measurements that use SPL stimulus calibrations less reliable at frequencies near 4 kHz and above. Studies have shown that FPL calibrations improve the reliability of DPOAE measurements over the standard SPL calibration (Maxim et al., 2019) and the results of this study suggest that use of FPL/EPL may help to expand the clinical use of DPOAEs to estimate hearing thresholds. With stimuli calibrated in FPL, which already reduces the variance between tests, measuring the DPOAE in EPL still improves the correlation between audiometric thresholds and DPOAE amplitudes. Handling calibration of not only stimulus levels but the level of the emission appropriately further reduced undesirable measurement noise to improve the diagnostic utility of DPOAEs.

The present study used swept tones rather than the clinical standard of testing discrete frequencies. This allows for a complete assessment across the frequency range with little additional time burden. Though swept tone DPOAEs are used in research applications (e.g., Abdala et al., 2015), their analysis is more complex, and norms have not been developed, which limits their current utility in clinical settings. At present, there is also no standardized protocol for swept otoacoustic emissions and their analysis, but adjustments to the least-squares fitting parameters can affect the resulting calculation of DPOAE amplitudes. Accordingly, discrete tone DPOAEs are likely to be more readily interpretable in clinical settings until more user-friendly analysis protocols are developed. However, the FPL and EPL calibration techniques are not exclusive to swept DPOAEs and are beneficial regardless of stimulus paradigm.

### B. Other Sources of Variability

Even when controlling stimulus levels, DPOAE amplitudes were highly variable across individual participants with similar hearing thresholds. For example, participants with normal hearing in the extended high frequencies had DPOAE amplitudes ranging 30 dB. Extended high-frequency audiometric thresholds can also be affected by issues related to standing waves in the ear canal. Like the DPOAEs probe, sound levels reaching the eardrum are affected by insertion depth, so thresholds are more variable in the extended high-frequency range when using insert earphones (Lapsley Miller et al., 2018). However, the high-frequency headphones used for this study have been shown to have reasonable test-retest reliability. Frank (2001) found that 98% of thresholds obtained with Sennheiser HAD 200 headphones were within 10 dB of the thresholds on subsequent tests.

Though this range is accepted clinically, it is another source of noise that may obscure the true relationship between tests. Depth-compensated calibrations such as FPL can also be used for measurement of extended high-frequency audiometric thresholds, though test-retest of high frequency thresholds was still ∼10 for at least one study (Lee et al., 2012).

Accurate calibrations for both measures, separation of the distortion and reflection components of the DPOAE, and careful data collection are not likely to result in a perfect correlation between DPOAEs and behavioral thresholds. The audiogram provides a more functional assessment of perception while DPOAEs reflect cochlear function of the outer hair cells. Though cochlear dysfunction is likely to affect both, the pathways they assess differ. But the more the non-physiologic sources of noise that contaminate diagnostic tests are controlled, the greater confidence we can have that the results reflect the physiologic processes our measures were designed to assess. Diagnostic precision is key to early identification and treatment of hearing loss and, more broadly, will improve understanding of individual differences that matter for hearing outcomes.

### C. Relationship with Age

Multiple studies have shown that age is a strong predictor of hearing thresholds (Carcagno and Plack, 2020; Grant et al., 2022; Lee et al., 2012; Wang et al., 2021); our results show a similar relationship between age and DPOAE emission amplitudes. Although our cohort was chosen specifically to have normal hearing sensitivity through 8 kHz, nearly half of our subjects had extended high-frequency hearing loss, and the proportion of subjects with extended high-frequency loss grew with each decade of life. This subclinical hearing loss may have practical implications for speech-in-noise understanding and suggests that extended high-frequency audibility may be a potential confound for studies relating age to auditory perception (Lough and Plack, 2022).

### D. Potential Clinical Applications

Future work is needed to implement new calibration strategies like FPL into clinical systems. Translation to the audiology clinic and expanded use in research settings may rely on the development of accessible hardware and software. The equipment used for this study (ER-10X) is expensive and no longer commercially available, but alternative custom calibration tubes can be made (Scheperle et al., 2008). Other strategies have been suggested to improve calibration-related errors (Souza et al., 2014), and complex integrated pressure may be a more practical calibration method to implement than FPL (Norgaard and Bray, 2023). More studies will be needed to fully optimize calibration, data collection, and data analysis protocols to ensure robust estimates of DPOAEs can be collected efficiently.

If such calibrations can be implemented clinically, one valuable use-case would be ototoxicity monitoring programs. Measuring extended high-frequency hearing may allow for earlier identification of hearing loss, leading to better informed individualized treatment plans and clinical recommendations. For example, the American Academy of Audiology recommends that persons undergoing treatment with an ototoxic medication should receive hearing screenings before, during, and after treatment, encouraging the use of high frequency audiometry and OAEs to monitor for hearing loss (Durrant et al., 2009). A change in hearing and its effect on post-treatment quality of life can be considered in conjunction with the medical necessity of the treatment. Just as a change in audiometric thresholds may inform clinical decisions about treatment, decreases in DPOAE amplitude can indicate the need for medication adjustments to ensure hearing is preserved as much as possible.

The ability to estimate hearing thresholds from DPOAEs also opens the door to more efficient diagnostic testing of infants or other populations that are difficult to test behaviorally. Given the potential perceptual implications of extended high-frequency hearing loss and its high prevalence, expanded testing of those frequencies in the audiology clinic may prove beneficial.

## V. CONCLUSION

Reliable physiological measures in the extended high frequencies are important for identifying hearing loss and improved diagnostic tools are needed for greater personalization of treatment for hearing loss. Critical to optimizing these measures is reducing known sources of variability as much as possible. With FPL/EPL calibration methods, DPOAEs can serve as a valuable tool to assess and monitor extended high-frequency hearing as it correlates with behavioral hearing thresholds.

## ACKNOWLEDGMENTS

This work was supported by NIH R01DC015989 (HMB, ARHM, AW), NIH R01DC009838 (SNH), NIH T32DC016853 (SNH), and NIH F32DC021345 (SNH).

## AUTHOR DECLARATIONS

## Ethics Approval

This research was approved by the Purdue Institutional Review Board (#1609018209).

Informed consent was obtained from all participants.

## DATA AVAILABILITY

The data that support the findings of this study were originally collected as part of Bharadwaj et al., 2022 and can be obtained from https://github.com/haribharadwaj/CommunBiol_CrossSpecies_Synaptopathy. This data set is also permanently archived using Zenodo and can be obtained from https://doi.org/10.5281/zenodo.6672827.

